# Dissecting the neural focus of attention reveals distinct processes for spatial attention and object-based storage in visual working memory

**DOI:** 10.1101/347518

**Authors:** Nicole Hakim, Kirsten C. S. Adam, Eren Gunseli, Edward Awh, Edward K. Vogel

**Author notes:** **Correspondence to:** Nicole Hakim, University of Chicago, 940 E 57^th^ St, Chicago, IL 60637. **Data Availability:** Datasets for all experiments will become available online on Open Science Framework upon acceptance of the manuscript or reviewer request: https://osf.io/ws3j9/?view_only=40b7f0018df44d5a862b51601d4599ed.

## Abstract

Complex cognition relies on both online representations in working memory (WM) said to reside in the *focus of attention*, and passive offline representations of related information. Here, we dissect the focus of attention by showing that distinct neural signals index the online storage of objects and sustained spatial attention. We recorded EEG activity during two tasks that employed identical stimulus displays while the relative demands for object storage and spatial attention varied. We found distinct delay-period signatures for an attention task (which only required spatial attention) and WM task (which invoked both spatial attention and object storage). Although both tasks required active maintenance of spatial information, only the WM task elicited robust contralateral delay activity that was sensitive to mnemonic load. Thus, we argue that the focus of attention is maintained via a collaboration between distinct processes for covert spatial orienting and object-based storage.

## Introduction

Working memory (WM) facilitates the temporary maintenance of small amounts of information so that it can be manipulated or acted upon. Contemporary theories of WM have coalesced on variations of embedded process models (Cowan, 1999; Oberauer, 2002) in which performance in WM tasks depends upon memory mechanisms that represent information in two distinct states: an online, active state (“focus of attention”); and an offline, passive state (“silent WM”). The focus of attention generally refers to information that is currently “in mind”, whereas silent WM is information that was recently within the focus but can still be rapidly accessed. These memory states have been proposed to be implemented at the neural level through persistent neural firing for items within the focus (Curtis & D’Esposito, 2003) and via rapid synaptic plasticity that allows recently attended items to quickly be reinstated (Jonides, Lacey, & Nee, 2015; Lewis-Peacock, Drysdale, Oberauer, & Postle, 2012; Rose et al., 2016; Stokes, 2015; Wolff, Jochim, Akyurek, & Stokes, 2017).

Here, we seek to characterize the neural mechanisms supporting the focus of attention. Broad neuroscientific support for focus of attention-related activity has been observed in sustained neural firing in monkey electrophysiological studies (Buschman, Siegel, Roy, & Miller, 2011; Funahashi, Chafee, & Goldman-Rakic, 1993), uni- and multivariate measurements of BOLD in human fMRI studies (Cowan et al., 2011; Majerus et al., 2016; Todd & Marois, 2004; Xu & Chun, 2006), and sustained electrical and magnetic fluctuations in human EEG and MEG studies (van Dijk, van der Werf, Mazaheri, Medendorp, & Jensen, 2010; Vogel & Machizawa, 2004). Within EEG and MEG studies, two candidate measures are consistent with the focus of attention construct. The first is alpha power (8-12hz), which shows sustained modulations during the retention period and has been shown to contain precise spatial information about the remembered/attended stimulus (Foster, Bsales, Jaffe, & Awh, 2017; Foster, Sutterer, Serences, & Awh, 2016). Another candidate is the Contralateral Delay Activity (CDA), which is a sustained negativity over the hemisphere contralateral to the positions of to-be-remembered items. CDA amplitude is modulated by the number of items held in WM, reaches an asymptote once WM capacity is exhausted, dynamically tracks dropping information, and predicts individual differences in WM capacity (Unsworth, Fukuda, Awh, & Vogel, 2014; Vogel & Machizawa, 2004; Vogel, McCollough, & Machizawa, 2005; Williams & Woodman, 2013). A prevailing view of the CDA is that it tracks the number of task-relevant objects that are stored in WM (Balaban & Luria, 2017; Luria, Balaban, Awh, & Vogel, 2016).

While the literatures on the CDA and alpha power have largely developed independently, recent proposals claim that they reflect isomorphic measures of the focus of attention. Specifically, van Dijk, et al. (2010) argued that the CDA is an averaging artifact of trial-level alpha modulation, and, therefore, reflects attention to the spatial positions of the memoranda, rather than representations of items in WM. A similar proposal was made by Berggren & Eimer (2016), who found that when two arrays were presented sequentially in different hemifields, CDA amplitude tracked the positions of the most recently seen items (but also see: Feldmann-Wüstefeld, Vogel, & Awh (2018). Such spatial attention accounts make two broad, but untested assertions regarding neural measures of the focus of attention. First, that sustained EEG activity reflecting the focus of attention exclusively represents the current regions of attended space, rather than the online maintenance of the items that occupy those regions of space (Berggren & Eimer, 2016). Second, that such neural measures amount to a monolithic “focus of attention,” rather than a collection of distinct, but overlapping mechanisms that together comprise the focus of attention.

Here, we provide evidence that the focus of attention in WM is not a monolithic construct, but rather, involves at least two neurally separable processes: (1) attention to regions in space (2) representations of objects that occupy the attended regions (i.e., object files). Alpha activity, but not the CDA, tracked attention to relevant spatial positions. Conversely, when participants stored object representations, lateralized alpha activity that tracked the attended positions was accompanied by robust, load-sensitive CDA. These results suggest the neural focus of attention can be dissected into at least two complementary, but distinct facets of activity: a map of prioritized space and online representations of objects.

## Materials & Methods

### Experimental Design

Our broad strategy was to compare delay period activity across two tasks that employed physically identical displays but distinct cognitive requirements. We designed distinct “Attention” and “WM” tasks to disentangle the neural correlates of hypothesized sub-components of the focus of attention. Both tasks are known to recruit sustained spatial attention (i.e. representation of a spatial priority map), but only the WM task invoked online storage of items (i.e. representation of the objects which occupied the attended locations). For all experiments, participants completed both a WM task and an attention task, and the sequence of physical stimuli was identical for both tasks; the attention and WM tasks differed only in the instructions given to participants and in the response mapping to keys. In Experiment 1, the WM task required that participants remember the color of the items in the sample array, whereas the attention condition required participants to direct spatial attention towards the locations of the items in the sample array (item color was irrelevant). Although highly similar, one key difference between the tasks in Experiment 1 was that participants were required to remember non-spatial features only in the WM task. To test whether the requirement to remember non-spatial features was responsible for our findings in Experiment 1, we employed even more similar tasks in Experiment 2. The WM task required that participants *store the spatial positions of items* in the sample array, and the attention task required that participants *covertly attend spatial positions* in anticipation of rare targets during the delay.

### Participants

Experimental procedures were approved by the University of Chicago Institutional Review Board. All participants gave informed consent and were compensated for their participation with cash payment ($15 per hour); participants reported normal color vision and normal or corrected-to-normal visual acuity. Participants were recruited from the University of Chicago and surrounding community. For each sub-experiment (e.g., Exp. 1a), we set a minimum sample size of 20 subjects (after attrition and artifact rejection). This minimum sample size was chosen to ensure that we would be able to robustly detect set-size dependent delay activity. Prior work employing sample sizes of 10 to 20 subjects per experiment can robustly detect set-size dependent CDA (Vogel & Machizawa, 2004; Vogel et al., 2005), and differences in CDA amplitude between novel experimental conditions (Balaban & Luria, 2017). We chose a minimum sample size toward the upper end of this conventional range.

A total of 63 and 54 participants were run in Experiments 1 and 2, respectively. Due to a technical error, EEG activity was not recorded for 3 participants in Experiment 1. In addition, data from some participants was excluded because of excessive EEG artifacts (<120 trials remaining in any of the four experimental conditions) or poor behavioral performance. This left 48 subjects in Experiment 1 (28 in Experiment 1a, 20 in Experiment 1b), and 49 subjects in Experiment 2 (20 in Experiment 2a, 29 in Experiment 2b).

### EEG Acquisition

Participants were seated inside an electrically shielded chamber, with their heads resting on a padded chin-rest 74 cm from the monitor. We recorded EEG activity from 30 active Ag/AgCl electrodes (Brain Products actiCHamp, Munich, Germany) mounted in an elastic cap positioned according to the International 10-20 system [Fp1, Fp2, F7, F8, F3, F4, Fz, FC5, FC6, FC1, FC2, C3, C4, Cz, CP5, CP6, CP1, CP2, P7, P8, P3, P4, Pz, PO7, PO8, PO3, PO4, O1, O2, Oz]. Two additional electrodes were affixed with stickers to the left and right mastoids, and a ground electrode was placed in the elastic cap at position Fpz. Data were referenced online to the right mastoid and re-referenced offline to the algebraic average of the left and right mastoids. Incoming data were filtered [low cut-off = .01 Hz, high cut-off = 80 Hz, slope from low-to high-cutoff = 12 dB/octave] and recorded with a 500 Hz sampling rate. Impedance values were kept below 10 kΩ.

Eye movements and blinks were monitored using electrooculogram (EOG) activity and eye-tracking. We collected EOG data with 5 passive Ag/AgCl electrodes (2 vertical EOG electrodes placed above and below the right eye, 2 horizontal EOG electrodes placed ~1 cm from the outer canthi, and 1 ground electrode placed on the left cheek). We collected eye-tracking data using a desk-mounted EyeLink 1000 Plus eye-tracking camera (SR Research Ltd., Ontario, Canada) sampling at 1,000 Hz. Usable eye-tracking data were acquired for 25 out of 28 participants in Experiment 1a, 19 out of 20 participants in Experiment 1b, 17 out of 20 participants in Experiment 2a, and 29 out of 29 participants in Experiment 2b.

### Artifact rejection

Eye movements, blinks, blocking, drift, and muscle artifacts were first detected by applying automatic detection criteria. After automatic detection, trials were manually inspected to confirm that detection thresholds were working as expected. Subjects were excluded if they had fewer than 120 total trials remaining in any of the 4 conditions. In Experiment 1a, we rejected an average 25% of trials across all four conditions. This left us with an average of 282 trials in WM set size 2 condition, 275 trials in the WM set size 4 condition, 302 trials in the Attention set size 2 condition, and 302 trials in the Attention set size 4 condition. In Experiment 1b, we rejected an average of 32% of trials across all four conditions. This left us with an average of 291 trials in the WM set size 2 condition, 285 trials in the WM set size 4 condition, 320 trials in the attention set size 2 condition, and 320 trials in the attention set size 4 condition. In Experiment 2a, we rejected an average of 22% of trials across all four conditions. This left us with an average of 302 trials in the WM set size 2 condition, 301 trials in the WM set size 4 condition, 322 trials in the attention set size 2 condition, and 323 trials in the attention set size 4 condition. In Experiment 2b, we rejected an average of 27% of trials across all four conditions. This left us with an average of 283 trials in the WM set size 2 condition, 283 trials in the WM set size 4 condition, 298 trials in the attention set size 2 condition, and 295 trials in the attention set size 4 condition.

#### Eye movements

We used a sliding window step-function to check for eye movements in the HEOG and the eye-tracking gaze coordinates. For HEOG rejection, we used a split-half sliding window approach (window size = 100 ms, step size = 10 ms, threshold = 20 μV). We only used the HEOG rejection if the eye tracking data were bad for that trial epoch. We slid a 100 ms time window in steps of 10 ms from the beginning to the end of the trial. If the change in voltage from the first half to the second half of the window was greater than 20 μV, it was marked as an eye movement and rejected. For eye-tracking rejection, we applied a sliding window analysis to the x-gaze coordinates and y-gaze coordinates (window size = 100 ms, step size = 10 ms, threshold = 0.5° of visual angle).

#### Blinks

We used a sliding window step function to check for blinks in the VEOG (window size = 80 ms, step size = 10 ms, threshold = 30 μV). We checked the eye-tracking data for trial segments with missing data-points (no position data is recorded when the eye is closed).

#### Drift, muscle artifacts, and blocking

We checked for drift (e.g. skin potentials) by comparing the absolute change in voltage from the first quarter of the trial to the last quarter of the trial. If the change in voltage exceeded 100 μV, the trial was rejected for drift. In addition to slow drift, we checked for sudden step-like changes in voltage with a sliding window (window size = 100 ms, step size = 10 ms, threshold = 100 μV). We excluded trials for muscle artifacts if any electrode had peak-to-peak amplitude greater than 200 μV within a 15 ms time window. We excluded trials for blocking if any electrode had at least 30 time-points in any given 200-ms time window that were within 1μV of each other.

### Analysis of Horizontal Gaze Position

We rejected all trials that had eye movements greater than 0.5° of visual angle. Nevertheless, participants could still move their eyes within the 0.5° of visual angle threshold (e.g. microsaccades). To compare eye movements in the two tasks, we compared the horizontal gaze position recorded by the eye tracker. We were most concerned with horizontal eye movements, as these could contaminate our lateralized EEG measures. We drift-corrected gaze position data by subtracting the mean gaze position measured 200 ms before the pre-cue to achieve optimal sensitivity to changes in eye position (Cornelissen, Peters, & Palmer, 2002). We then took the mean change in gaze position (in degrees of visual angle) for left and right trials during same time-window that we used in the CDA analysis, 400 to 1450 ms after stimulus onset. Eye gaze values from left trials were sign-reversed so that left and right trials could be combined together. As such, positive values indicate eye movements toward the remembered side, and negative values indicate eye movements away from the remembered side. Importantly, not all participants had eye tracking with adequate quality to be included in this analysis. Therefore, only 25 participants from Experiment 1a, 19 participants from Experiment 1b, 17 participants from Experiment 2a and 27 participants from Experiment 2b were included in the analysis.

### Analysis of Pupil Dilation

As an additional metric of task difficulty, we compared task-evoked pupil dilation between the WM and attention tasks. Many studies have demonstrated that task-evoked pupil dilation correlates with cognitive load; the pupil dilates more when there are higher attentional and working memory demands (Beatty, 1982; Steinhauer & Hakerem, 1992). Since we were most interested in assessing the relative difficulty of the two tasks, we collapsed the data across set size within each task. For our analysis, pupil dilation data were baselined from 400 to 0 ms before the onset of the colored squares. Differences in pupil dilation between the WM and attention tasks (collapsed across set sizes) were calculated by comparing pupil size during the same time-window as is used in the CDA analysis (400 to 1450 ms after stimulus onset). Just as in the analysis of horizontal gaze position, not all participants had eye tracking that was good enough to be included in this analysis. The same participants were included in both the horizontal gaze position and pupil dilation analyses.

### Analysis of contralateral delay activity

EEG activity was baselined from 400 ms to 0 ms before the onset of the stimulus array. Trials containing targets for the attention task were excluded. Event-related potentials were calculated by averaging baselined activity at each electrode across all accurate trials within each condition (Set-Size 2 WM, Set-Size 4 WM, Set-Size 2 Attention, and Set-Size 4 Attention). We calculated amplitude of contralateral and ipsilateral activity for five posterior and parietal pairs of electrodes chosen a priori based on prior literature: O1/O2, PO3/PO4, PO7/PO8, P3/P4, and P7/P8. Statistical analyses were performed on data that was not filtered beyond the .01 – 80 Hz online data acquisition filter; we low-pass filtered data (30 Hz) for illustrative purposes in paper figures.

### Analysis of lateralized alpha power

EEG signal processing was performed in MATLAB 2015a (The MathWorks, Natick, MA). We band-pass filtered trial epochs in the alpha band (8-12 Hz) using a bandpass filter from the FieldTrip toolbox (Oostenveld, Fries, Maris, & Schoffelen, 2011; ‘ft_preproc_bandpass.m’) and then extracted instantaneous power by applying a Hilbert transform (‘hilbert.m’) to the filtered data. Trials containing targets for the attention task were excluded. We calculated alpha power for the same five posterior and parietal pairs of electrodes as CDA: O1/O2, PO3/PO4, PO7/PO8, P3/P4, and P7/P8.

### Stimuli & Procedures

Stimuli in all experiments were presented on a 24-inch LCD computer screen (BenQ XL2430T; 120 Hz refresh rate) on a Dell Optiplex 9020 computer. Participants were seated with a chin-rest 74 cm from the screen. Stimuli were presented on a gray background, and participants fixated a small black dot with a diameter of approximately 0.2 degrees of visual angle.

#### Experiment 1a

We ran two very similar versions of Experiment 1 (hereafter referred to as Experiments 1a and 1b). The stimuli and procedures for Experiments 1a and 1b were almost identical, with the exception of the differences noted in the “Experiment 1b” section. Each trial began with a blank inter-trial interval (750 ms), followed by a diamond cue (300 ms) indicating the relevant side of the screen (right or left). This diamond cue (0.65° maximum width, 0.65° maximum height) was centered 0.65° above the fixation dot and was half green (RGB = 74, 183, 72) and half pink (RGB = 183, 73, 177). Half of the participants were instructed to attend the green side and the other half were instructed to attend the pink side. After the cue, 2 or 4 colored squares (Exp. 1: 1.1° by 1.1°; Exp. 2: 1° by 1°) briefly appeared in each hemifield (150 ms) with a minimum of 2.10° (1.5 objects) between each item. The squares then disappeared for a 1,300 ms delay period. Squares could appear within a subset of the display subtending 3.1° to the left or right of fixation and 3.5° degrees above and below fixation. Colors for the squares were selected randomly from a set of 9 possible colors (Red = 255 0 0; Green = 0 255 0; Blue = 0 0 255; Yellow = 255 255 0; Magenta = 255 0 255; Cyan = 0 255 255; Orange = 255 128 0; White = 255 255 255; Black = 1 1 1). Colors were chosen without replacement within each hemifield, and colors could be repeated across, but not within, hemifields. On 10% of trials, two small, black lines (0.02° wide, 0.4° long) appeared (66.7 ms), one at the location of a colored square in the cued hemifield, and one at the location of a colored square in the uncued hemifield. The lines could appear at any point during the delay period from 100 to 1200 ms after the offset of the stimuli. Each line could be tilted 31.3 degrees to the left or 31.3 degrees to the right. At test, a probe display appeared until response, consisting of 1 colored square in each hemifield.

#### Experiment 1b

All stimuli and procedures were the same as Experiment 1a with the following exceptions. On 10% of trials, two small, black lines appeared, in the cued hemifield. One line appeared at the location of a colored square. The other line appeared in an unoccupied location in the cued hemifield with a minimum of 2.10° (1.5 objects) from the locations of the memory array items.

#### Experiment 2a

Stimuli were similar to Experiment 1b with the following exceptions. Participants were presented with 2 or 4 black circles (0.611° diameter; RGB= 1 1 1) in each hemifield with a minimum of 1.53° degrees (1.5 objects) between each item. These circles and distractor line could appear within a subset of the display subtending 2.44 degrees to the left or right of fixation and 3.06 degrees above and below fixation. On target-present trials, the two small lines that were presented briefly during the retention interval were 0.04° wide and 0.76° long. Both were presented in the attended hemifield. One was at the location of a colored square, and the other was in an unoccupied location that was a minimum of 1.5 objects away from any other memory location.

#### Experiment 2b

Stimuli were similar to Experiment 2a with the following exceptions. Participants were presented with 2 or 4 colored circles (0.84° diameter) in each hemifield with a minimum of 2.10° (1.5 objects) between each item. Colors for the circles were randomly selected without replacement within each hemifield from a set of 10 possible colors (Red = 255 0 0; Green = 0 255 0; Blue = 0 0 255; Yellow = 255 255 0; Magenta = 255 0 255; Cyan = 0 255 255; Orange = 255 128 0; Brown = 102 51 0; White = 255 255 255; Black = 1 1 1). On target-present trials, the two small lines that were presented briefly during the retention interval were 0.04° wide and 0.99° long. Both were presented in the attended hemifield. One was at the location of a colored square, and the other was in an unoccupied location that was a minimum of 1.5 objects away from any other memory location.

Participants in all experiments completed a WM and an attention task (Figure 1). Within each experiment, the sequence of physical stimuli was identical for both tasks. Differences in procedures between the experiments are described below. The attention and WM tasks differed only in the instructions given to participants and in the keys used to respond. Task order (attention first or WM first) and relevant cue color (pink or green) were counterbalanced across participants. Participants completed 20 blocks of 80 trials each (1,600 trials total, 400 per condition).

**Figure 1.**
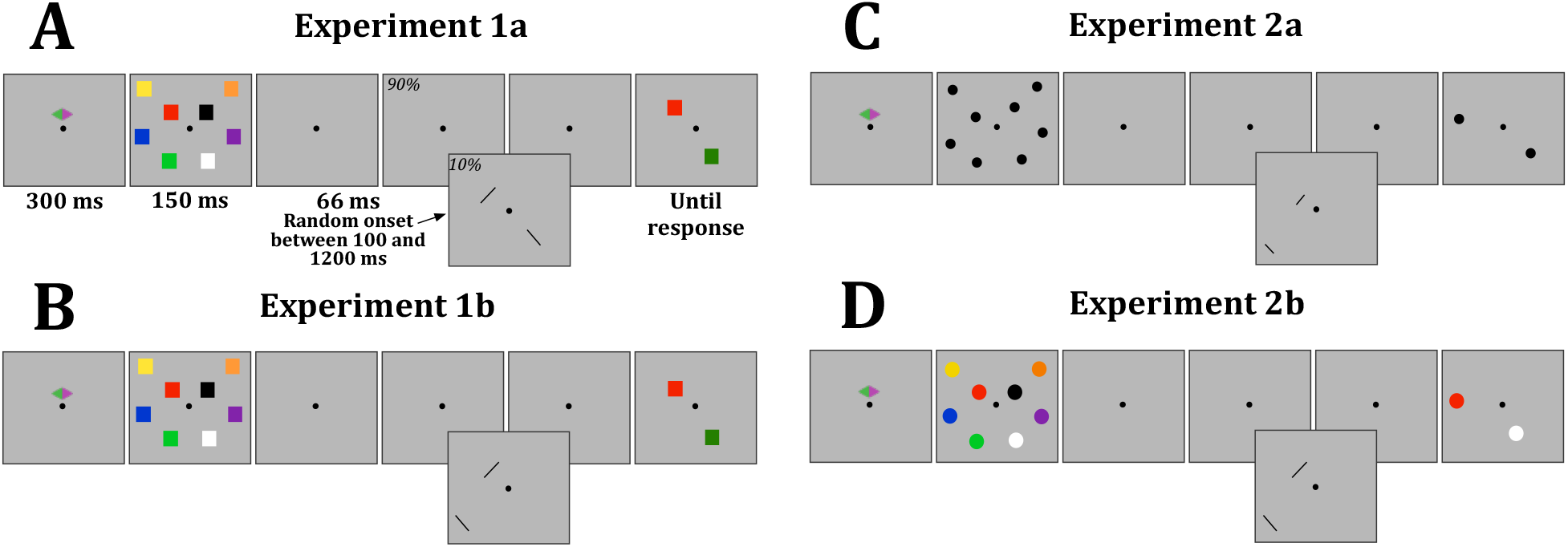
Working memory and attention tasks for all experiments. At the start of each trial, a cue appeared on the screen for 300 ms, which cued participants to attend one side of the screen. Then, an array of 2 or 4 colored squares (Exp. 1a & 1b) or circles (Exp. 2a & 2b) briefly appeared (150 ms). On 10% of trials, during the blank retention interval (1300 ms), two small lines appeared for 66 ms between 100 and 1200 ms after memory array offset. In Experiment 1a, one line appeared in each hemifield. In all other experiments, both lines appeared in the same hemifield, one in an attended location and one in an unattended location. After the retention interval, a response screen appeared with one square (Exp. 1a & 1b) or circle (Exp. 2a & 2b) in each hemifield. In the WM task, to respond, participants reported whether the square (Exp. 1a & 1b) or circle (Exp. 2a & 2b) that reappeared in the attended hemifield was the same color (Exp. 1a & 1b) or in the same location (Exp. 2a & 2b). Participants pressed “z” if it was the same color (Exp. 1a & 1b) or location (Exp. 2a & 2b) and “/?” if it was different. In the Attention task, if a line was not present during the delay period, participants pressed “spacebar.” If a line was present during the delay, participants had to report the orientation of the line that appeared in one of the cued locations. If the line was tilted left, participants pressed “z” and if it was tilted right, participants pressed “/?.” The response screen remained visible until a response was made.

#### Working Memory Task

In Experiments 1a and 1b, participants were instructed to remember the colors of the presented squares in the cued hemifield and to ignore the lines that might flash during the middle of the delay period. At test, participants were asked to identify whether the color presented at the relevant probed location was the same as the color held in mind (same trial) or different (change trial). The colors changed on 50% of trials. Participants pressed the “z” key to indicate the response “same” and pressed the “/” key to indicate “different”. For Experiment 2a and 2b, the procedures for the WM task were very similar to the procedure from Experiment 1a and 1b, except that participants were asked to identify whether the location of the presented circle in the attended hemifield was in the same or different location as any of the original circles.

#### Attention Task

Procedures and instructions for the attention task were identical in all experiments. Only the visual stimuli differed, so as to match the visual stimuli presented in the WM task. Participants were instructed to maintain their attention at the locations of the presented squares in the cued hemifield in order to identify the orientation of a small line that appeared at one of the attended locations on 10% of trials. Participants were instructed to press the “z” key if the line appeared and was tilted left, and the “/” key if the line appeared and was tilted right. On 90% of trials, no line was presented, and participants were instructed to press the “space” key to indicate that there was no target present. The physical stimulus displays were identical to the memory task; thus, one colored square appeared in each hemifield at the end of the attention trials. Participants were told that the appearance of the test display indicated that it was time to respond, and that the location and the color of the squares were irrelevant to the task.

Stimuli and procedures in Experiment 1a differed from Experiments 1b, 2a, and 2b only for the target-present trials (10% of trials). Specifically, in Experiments 1b, 2a, and 2b during target-present trials, we presented both a relevant and an irrelevant line within the cued hemifield. One line always appeared at the same location as one of the colored squares; the second line appeared at a foil location where no colored square had been presented (a minimum distance of 1.5 items, width from any of the colored squares, locations). Thus, the participants were required to maintain their attention at precise locations within the relevant hemifield so that they knew which line to report. We reasoned that the inclusion of an irrelevant item in the cued hemifield in Experiment 1b would encourage subjects to orient attention more precisely. However, subsequent analyses revealed no main effect or interactions associated with the changes in procedure between Experiment 1a and 1b. Therefore, data were collapsed across these two versions of the task.

We would additionally like to note that in both the memory and attention tasks, the circles/squares in the sample array were always at least 1.5 objects apart from each other. In the Attention task, the rare target probes appeared at the location of one of the original squares, while the distractor probe appeared in an uncued location that was at least 1.5 objects away from any of the attended locations. Thus, the Attention task required subjects to make the same spatial discrimination that subjects had to make in the memory task in order to relate the test probe to the proper item from the memory array. In other words, the positions of the sample items had to be maintained equally precisely in the memory and attention tasks. This is most clear for Experiment 2, in which space was the sole relevant attribute for the memory task.

## Results

We aggregated data across Experiments 1 and 2 (n=97) to provide the most power for understanding the distinctions between the WM and Attention tasks. While the aggregate results mirrored those of the individual experiments (see Supplemental Materials for a study-by-study analysis), the data taken together provide a clear demonstration of the essential empirical patterns. In this aggregate analysis, we focus on CDA, alpha power, and pupil size. For analyses of behavior and eye position for each study, see Supplemental Materials.

### Preliminary analysis of the effect of Experiment

In a preliminary analysis, we examined whether the small variations in task design between Experiments 1 and 2 had an effect on the observed results. For this purpose, we ran repeated-measures ANOVAs for each analysis (i.e., CDA, alpha power, and pupil size) with the within-subjects factors Task (WM, Attention) and Set Size (2, 4 items) and the between-subjects factor Experiment (1, 2). For all analyses, there was no main effect of Experiment, *p*>=.16. Therefore, it was justified to collapse data across the all experiments.

For the horizontal gaze position and the lateralized alpha analyses, none of the factors significantly interacted with Experiment, *p*>=.19. However, for the pupil dilation analysis, there was a significant interaction of Task and Experiment, *F*(1,86)=10.76, *p*=.002, η_p_^2^ =.11. This significant interaction is explained by greater pupil dilation in the Attention task than in the WM task in Experiment 2, but not in Experiment 1.

For the CDA analysis, there was a significant 3-way interaction of Laterality, Set Size and Experiment, *F*(1,95)=6.73, *p*=.01, η_p_^2^ =.07. To further delineate this three-way interaction, we ran follow-up ANOVAs with the factors Laterality (contra, ipsi) and Set Size (2, 4 items) for Experiment 1 and Experiment 2 separately. These follow-up analyses revealed that there was a significant interaction of Laterality and Set Size for Experiment 2, *F*(1,48)=16.88, *p*<.001, η_p_^2^ =.26, but not for Experiment 1, *F*(1,47)=.08, *p*=.78, η_p_^2^ =.002.

**Figure 2.**
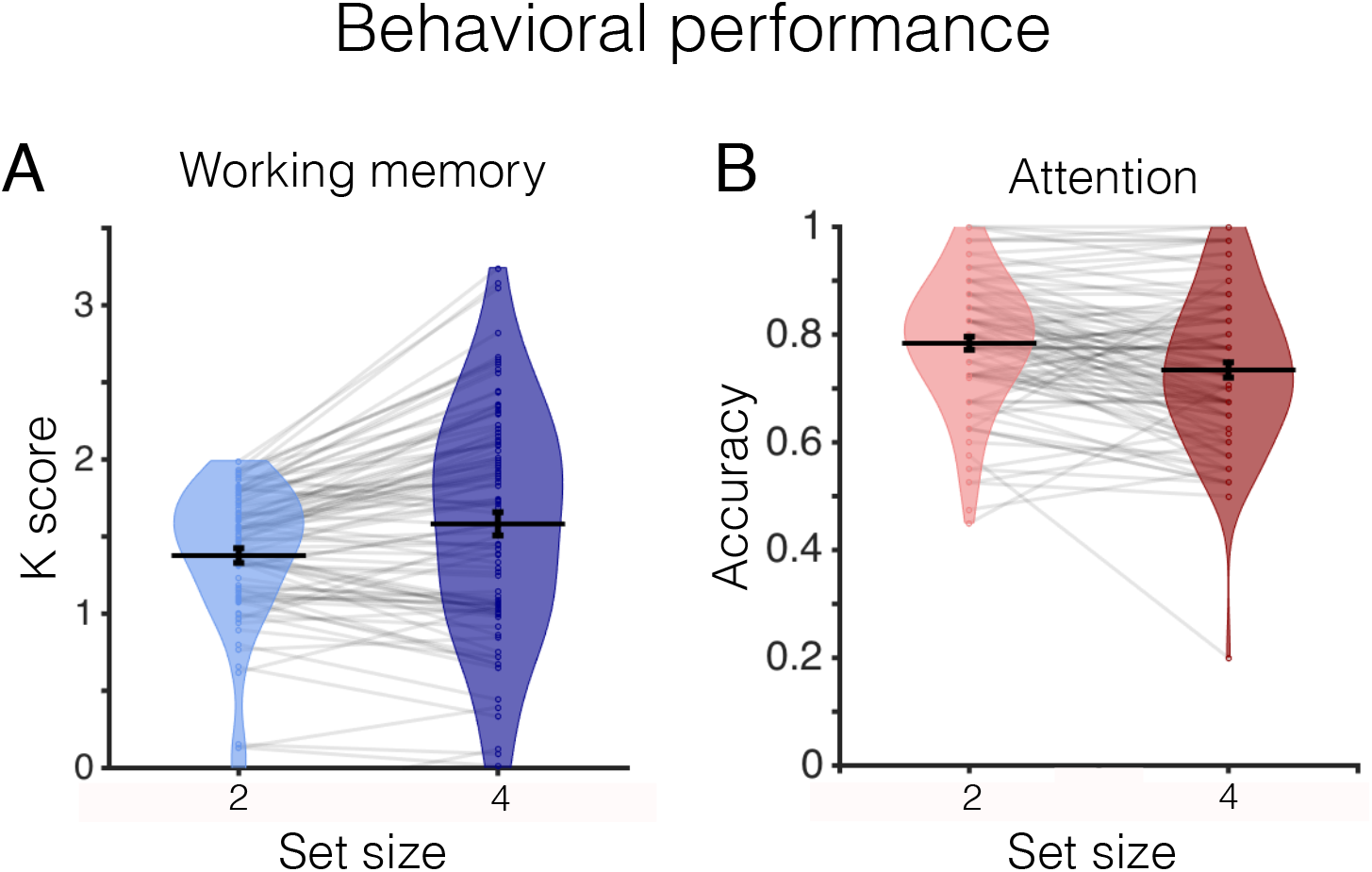
Behavioral performance. (A) Average K score in the WM task for all experiment. The distribution of K scores for all participants is represented by the violin plot. Dots and light gray lines represent one participant’s performance. (B) Average accuracy in the Attention tasks for all experiment.

### Contralateral Delay Activity

Using all data from Experiments 1 and 2 together, we ran a repeated-measures ANOVA with the factors Task (WM, Attention) and Set Size (2, 4 items). This analysis revealed significant main effects of Laterality, *F*(1,96)=74.41, *p*<.001, η_p_^2^ =.44, and Set Size, *F*(1,96)=33.27, *p*<.001, η_p_^2^ =.26. Because these main effects were collapsed across Task, they are not informative for our central question of how storage-related neural signals differ across tasks. Thus, the first important finding was a significant 2-way interaction between Laterality and Task, *F*(1,96)=81.27, *p*<.001, η_p_^2^ =.46, that reflected a greater laterality effect in the WM than in the Attention task. To confirm this impression, we ran a follow-up 2-way paired-samples t-test that compared contralateral to ipsilateral activity separately for the WM and Attention tasks and each set size (e.g. WM ss2, WM ss4, ATT ss2, ATT ss4). This analysis revealed that the CDA was significantly more lateralized in the WM (Set Size 2: M =-.38, SD=.44; Set Size 4: M=-.54, SD=.54) than in the Attention (Set Size 2: M=-.09, SD=.34; Set Size 4: M=-.10, SD=.37) task for both set sizes (Set Size 2: *t*(98)=-6.71, *p*<.001; Set Size 4: *t*(98)=-8.57, *p*<.001). We note, however, that there was reliable lateralized activity for both tasks, *p*<=.007.

Another key finding was that the number of items in the sample array had a selective impact on CDA in the WM task, while CDA in the Attention task showed no reliable effect. This impression was verified by a reliable triple interaction between Task, Laterality and Set Size, *F*(1,96)=8.75, *p*=.004, η_p_^2^ =.08. As Figure 3 shows, CDA was set size dependent in the WM task, but not in the Attention task. To characterize the triple interaction, we ran separate follow-up repeated-measures ANOVAs for each task (WM and Attention) with the factors Laterality (contra, ipsi) and Set Size (2, 4 items). This analysis revealed that there was a significant interaction of Laterality and Set Size for the WM task, *F*(1,96)=14.39, *p*<.001, η_p_^2^ =.13, but not the Attention task, *F*(1,96)=.07, *p*=.79, η_p_^2^ =.001. Thus, while data from the WM task showed that CDA amplitude was larger for set size 4 (M=-.55, SD=.54) than set size 2 (M=-.39, SD=.44), data from the Attention task showed no evidence of a difference in CDA amplitude between Set Size 2 (M=-.10, SD=.34) and Set Size 4 (M=-.11, SD=.37).

**Figure 3.**
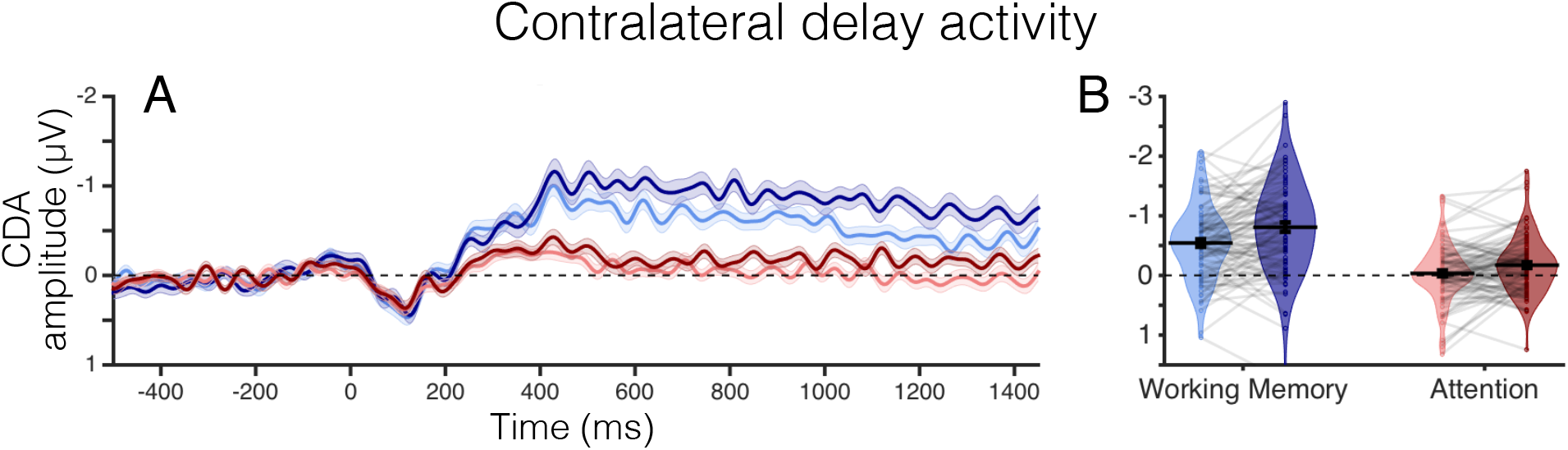
Contralateral delay activity. (A) CDA amplitude (μV) over time for all experiments combined. Timepoint zero marks the onset of the memory array, and timepoint 1450 marks the onset of the response array. (B) Average CDA amplitude (μV) for all experiments during the time window of interest, 400 to 1450 ms. The distribution of CDA amplitudes for all participants is represented by the violin plot. Dots and light gray lines represent one participant’s CDA amplitude.

To summarize, the aggregate analysis showed that CDA was substantially stronger in the WM than in the Attention task. Moreover, CDA tracked the increase in mnemonic load from two to four items, while the CDA signal in the Attention task – in addition to being over four times smaller than in the WM task – showed no effect of mnemonic load at all, a defining feature of the CDA. This core result motivates our conclusion that CDA is directly linked with the online maintenance of object representations in WM, and not the deployment of attention to the positions of the sample items.

### Lateralized Alpha Power

As Figure 4 shows, we observed the typical suppression of alpha power contralateral to the relevant hemifield in both tasks, though it was larger in the WM task. We confirmed these impressions with a repeated-measures ANOVA on the average alpha power with the factors Laterality (contra, ipsi), Task (attention, WM) and Set Size (2, 4 items). This analysis revealed a significant main effect of Laterality, *F*(1,96)=45.57, *p*<.001, η_p_^2^ =.32, and a significant interaction between Laterality and Set Size, *F*(1,96)=9.75, *p*=.002, η_p_^2^ =.09. Paired t-tests confirmed that this interaction reflects a stronger lateralization of alpha power in the set size 2 condition (M=-12.24, SD=17.17) than in the set size 4 condition (M=-10.04, SD=16.06) (*t*(96)=-3.123, *p*=.002). Thus, the strength of lateralized alpha activity varied with the number of stored or attended positions in both the WM and Attention tasks. Critically, however, the effect of set size on lateralized alpha power was in the opposite direction from the effect we observed with CDA. CDA was stronger for set size 4 than for set size 2 whereas alpha lateralization was stronger for set size 2 than for set size 4. These findings support the hypothesis that CDA and alpha activity reflect distinct aspects of online storage in visual WM.

**Figure 4.**
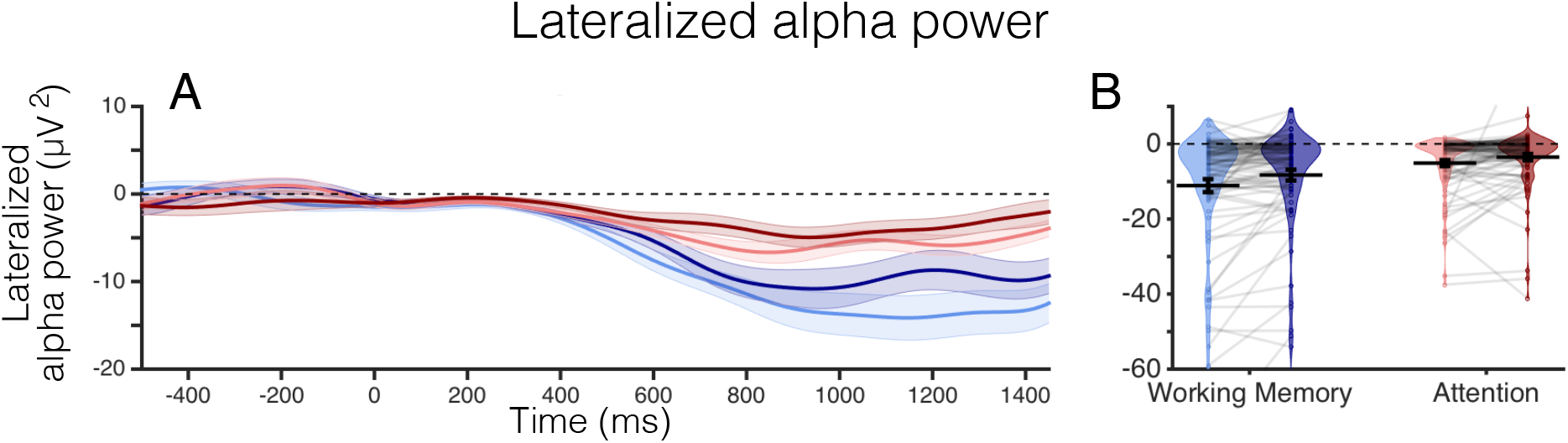
Lateralized Alpha Power. (A] Lateralized alpha power (μV^2^) over time collapsed across all experiments. Timepoint zero marks the onset of the memory array, and timepoint 1450 marks the onset of the response array. (B) Average lateralized alpha power (μV^2^) for all experiments during the time window of interest, 400 to 1450 ms. The distribution of lateralized alpha power for all participants is represented by the violin plot. Dots and light gray lines represent one participant’s alpha power.

Our analysis also revealed a significant interaction between Laterality and Task, *F*(1,96)=27.22, *p*<.001, η_p_^2^ =.22, that reflected the greater lateralization of alpha power in the WM than in the Attention task. This impression was confirmed with a two-way paired samples t-test that revealed a significant difference in alpha power lateralization between the WM (M=-7.79, SD=11.61) and Attention (M=-3.35, SD=5.69) tasks, *t*(96>-5.22, *p*<.001. Critically, both tasks showed clear evidence of lateralized alpha power in both set sizes (*p*<.001 for all conditions), confirming that covert attention was deployed to the position of sample items in a sustained fashion in both tasks. The greater lateralization of alpha power in the WM than Attention task is a robust empirical pattern that is present in both experiments and in the aggregate analysis. Though we did not expect this pattern *a priori*, this reliable difference in alpha lateralization between the two tasks may reflect a direct influence of online object representations on the deployment of spatial attention.

### Pupil Dilation

We argue that the WM task encouraged online storage of object representations while the Attention task did not. Thus, the restriction of load-dependent CDA to the WM task could reflect a direct link between the CDA and item storage in WM. A clear alternative hypothesis, however, is that the WM task may differ from the Attention task in terms of the intensity or effort applied to the task rather than the specific cognitive operations that were invoked. While accuracy was similar (and off of ceiling) in the two tasks, this does not provide strong evidence for equivalent effort. Fortunately, pupil dilation measurements have been shown to provide a sensitive index of cognitive effort and arousal when bottom-up stimulus factors are controlled. Thus, we ran a two-way paired-samples t-test to examine whether pupil size differed during the time window in which CDA was measured. This analysis revealed a greater level of pupil dilation in the Attention task (M= .63, SD=1.89) compared to the WM task (M= .29, SD=2.20), *t*(87)=-3.13, *p*=.002, suggesting that the Attention task recruited greater levels of cognitive effort. Thus, our finding that CDA was far larger in the WM task cannot be explained by increased effort in the WM task. Indeed, pupil analysis of the aggregated data suggests that the WM task was the easier of the two. These findings argue for a difference in the nature of the cognitive operations evoked by the WM and Attention tasks, rather than in the degree to which similar operations were carried out.

**Figure 5.**
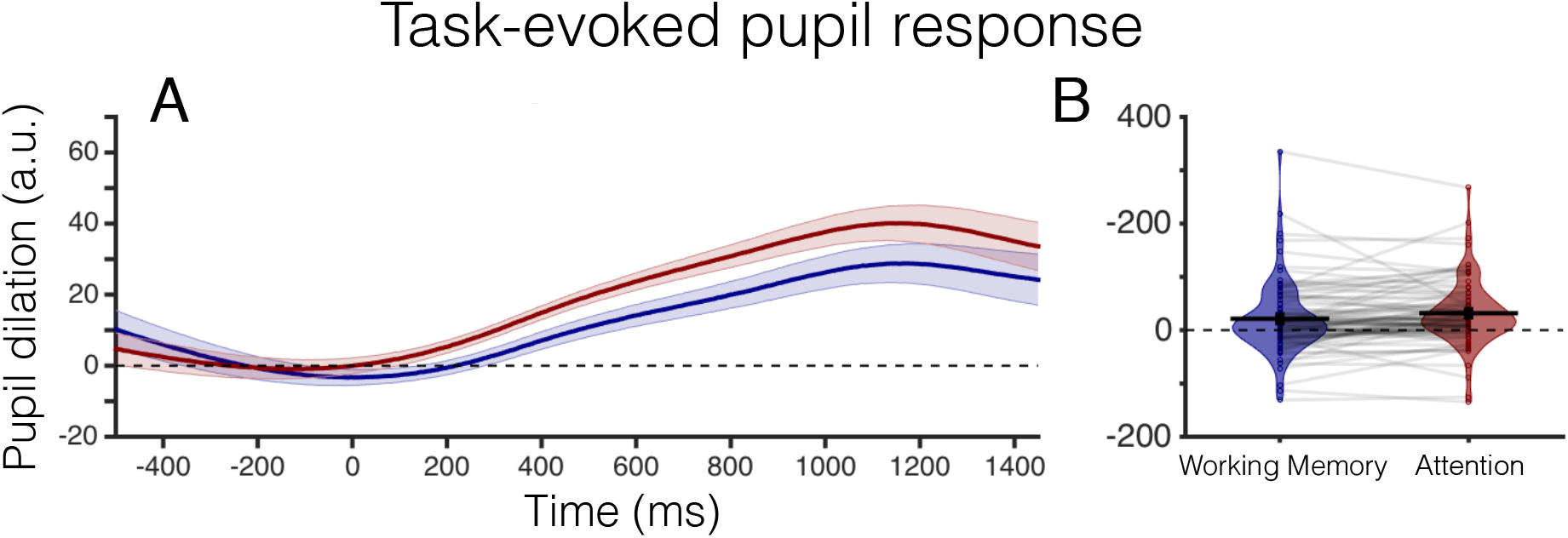
Task evoked pupil response. (A) Pupil dilation (arbitrary units) over time for all experiments combined. Timepoint zero marks the onset of the memory array, and timepoint 1450 marks the onset of the response array. (B) Average pupil dilation (arbitrary units) for all experiments during the time window of interest, 400 to 1450 ms. The distribution of pupil dilation for all participants is represented by the violin plot. Dots and light gray lines represent each participant’s pupil dilation.

## Summary

With 97 subjects, our aggregate analysis provided strong statistical power for documenting how neural activity differed between the WM and Attention tasks. CDA was more than four times larger in the WM task than in the Attention task. Moreover, CDA in the WM task clearly tracked changes in mnemonic load whereas CDA in the Attention task showed no evidence of load sensitivity. Thus, given that these tasks employed identical stimulus displays, we conclude that CDA may be directly tied to the unique *object representation* requirements in the WM task and not covert attentional orienting to the sample array positions.

The WM and Attention tasks differ in terms of object storage, but past work suggests that both WM and Attention tasks may call upon a common spatial attention process that elicits orderly changes in the scalp topography of alpha power. In addition to past studies showing the broad involvement of alpha activity across a wide range of attention and memory paradigms (Canolty & Knight, 2010; Fries, 2005; Klimesch, 2012), more recent work has also established that the topography of alpha activity on the scalp can be used to precisely track the locus of covert attention (Foster, Sutterer, Serences, Vogel, & Awh, 2017; Rihs, Michel, & Thut, 2007) and locations stored in WM (Foster, Bsales, et al., 2017; Foster et al., 2016). In line with this work, there was clear evidence from both the WM and Attention tasks that alpha power in posterior electrodes was reduced contralateral to the sample array. Importantly, the aggregate analysis also had enough power to reveal a reliable effect of set size on the strength of alpha lateralization, such that greater lateralization was observed in the set size 2 condition compared to the set size 4 condition. This effect should be interpreted with caution, however, as some previous research has not found an effect of set size on lateralization (Fukuda, Mance, & Vogel, 2015), while others have found greater lateralization for larger set sizes (Sauseng et al., 2009). Nevertheless, the fact that CDA showed the opposite pattern, with higher CDA for the larger set size, highlights the possibility that these two neural signals (measured from within the same set of electrodes) index distinct aspects of maintenance within the focus of attention.

There were differences in probe probability between the attention and working memory tasks. The working memory task required participants to make an attentionally demanding response after every trial. However, in the attention task, participants only had to make an attentionally demanding orientation discriminate on 10% of trials. On the remaining 90% of trials, participants had to press space, indicating that no lines were present. To our knowledge, the influence of probe probability on CDA amplitude has not been investigated. Therefore, the difference in CDA amplitude in the two tasks could be affected by probe probability. We think this alternate explanation is unlikely, as a large body of research has shown that the CDA tracks information stored in working memory across a wide range of response modalities, including two-alternative change detection (Vogel & Machizawa, 2004), whole report of discrete colors (Adam, Robison, & Vogel, 2018), and tracking of dynamic displays (Balaban & Luria, 2017; Drew & Vogel, 2008).

We additionally examined whether the observed differences in neural activity were a consequence of differential effort or arousal in the two tasks. Because the stimulus displays were identical, we were able to use task-evoked pupil dilation to obtain a sensitive metric of cognitive effort and arousal. The aggregate analysis revealed that the Attention task elicited reliably larger pupil size than the WM task, suggesting that the Attention task elicited greater effort. In line with this conclusion, we also note that while behavioral data from the Attention task showed that monitoring four locations was more difficult than monitoring two locations, CDA in the Attention task was unaffected by set size. Therefore, our findings argue strongly against the hypothesis that stronger delay period signals in the WM task were a consequence of greater cognitive effort.

## Discussion

The focus of attention refers to the small set of mental representations that can be held in an *online* or readily accessible state. Motivated by its central role in intelligent behaviors, there has been a longstanding effort to elucidate the neural signals that track the contents of this internally attended information. This body of work has tended to treat the focus as a monolithic entity, and has claimed that internal attentional processes influence selection and maintenance of cognitive representations in the absence of sensory input (Chun, 2011; Chun, Golomb, & Turk-browne, 2011). This idea has been supported by neural evidence that has found that sustained working memory representations in the brain occur in the same regions as perceptual representations, which are inherently modulated by attentional mechanisms (Postle, 2006). However, in this study, we extend the growing evidence that the focus of attention may be implemented via multiple component processes playing distinct functional roles: one that represents currently prioritized space (alpha); and another that reflects item storage within the focus of attention (CDA). This proposal converges with other findings that suggest a dissociation between spatial attention and WM storage (Tas et al., 2016; Sheremata et al 2018).

### Contralateral delay activity and lateralized alpha power: Distinct components of the focus of attention

van Dijk et al. (2010) proposed that asymmetric modulations of alpha power at the trial-level can generate a CDA-like negative slow wave in an event-related average. However, there is growing evidence that these two measures can be clearly dissociated. For example, Fukuda, Kang, & Woodman (2016) used a lateralized change detection task where they cued participants to one side of the screen, but had a longer than normal (1,000 ms) SOA between the cue and the memory array. During this blank cue period, participants knew which hemifield would contain memory items, but no items had yet appeared. During this time, there was robust alpha power lateralization but no CDA. However, after the memory array appeared, the CDA and alpha power lateralization appeared in concert during the memory maintenance period (1,000 ms). These results suggest lateralized alpha power, and thus attention, can be shifted to empty space, but that the CDA necessitates object storage (see also Fukuda, Mance & Vogel, 2015). A similar dissociation between attentional deployment to objects and to spatial location has been found with the anticipatory N2PC, a component which is related the CDA (Woodman, Arita, & Luck, 2009).

### Contralateral delay activity as an index of item-based storage in working memory

What was the critical difference between the WM and Attention tasks? Despite the fact that they employed identical stimulus displays, the amplitude of the CDA was more than four times larger in the WM than in the Attention task, and only the WM task elicited load-dependent CDA. Both tasks elicited covert orienting to the positions of the items in the sample array, as shown by sustained lateralized alpha power modulations. Moreover, despite distinct monikers, *both* tasks required the sustained maintenance of spatial information across a blank delay. This storage requirement is obvious for the WM change detection task. But even for the Attention task, subjects must have maintained the cued positions so that they could distinguish targets from lures. Indeed, in all experiments, change detection in the WM task required precisely the same spatial discriminations as did target identification in the Attention task. Thus, we propose that the critical difference between the WM and Attention tasks was that the WM task encouraged the continued *representation of the items* in the sample array, while in the Attention task participants directed spatial attention to those positions without maintaining the items themselves.

### Contralateral delay activity as a neural index of object file maintenance

Our interpretation of the CDA as an index of continued representations of object files critically hinges on a distinction between the maintenance of items in working memory and the maintenance of spatial information without an accompanying item representation. While some may view this as provocative, recent work has shown dissociable patterns of activity in parietal lobe between WM and spatial attention demands (Sheremata, Somers, & Shomstein, 2018). Additionally, we note that there is a longstanding precedent for a distinction between the representation of an object and the representation of the features or identifying labels associated with that object. Kahneman, Treisman, & Gibbs (1992) elucidated this idea with the *object file* construct which proposes two separable stages of processing. The first involves the parsing of the scene into a set of individuated items that are indexed based on their spatial and temporal coordinates. Subsequently, the specific feature values (e.g., color and orientation) are processed and incorporated into the associated object file. Thus, object files anchor the episodic representation in a specific time and place, and are distinct from the specific feature values that are bound together by virtue of an object file. In the present context, an intriguing possibility is that CDA indexes the maintenance of object files in WM. This proposal is consistent with recent work showing that the CDA is sensitive to objecthood cues (Balaban & Luria, 2016) and tracks the number of encoded objects, not the number of features within objects (Luria & Vogel, 2011). Thus, even though the Attention task required the sustained maintenance of location information, CDA was minimal or absent (and insensitive to mnemonic load) because the task did not encourage the maintenance of the object files that were created during the encoding of the sample array. Finally, while object files have been argued to mediate the binding of multiple features within an object, we note that this does not preclude the operation of object files for single-feature objects (Kahneman et al., 1992), such as those required by the spatial WM task of Experiment 2. Thus, we propose that the CDA may provide a neural index of object file maintenance.

### Open question on the impact of “object files” on the allocation of spatial attention

In this series of experiments, lateralized alpha power was a useful tool to illustrate that participants sustained their attention to the cued side even when the CDA was completely absent (Exp 1). However, we also observed a main effect of our task manipulation on lateralized alpha power. When task demands required participants to encode object representations, alpha power was significantly more lateralized than when they only had to sustain their attention to empty space. Though we did not predict this pattern *a priori*, it was reliable in both experiments. This suggests that, like the CDA, lateralized alpha power respects the dissociation between forming object representations and maintaining a spatial priority map. One possible interpretation of this effect is that object representations serve as “anchors” for the allocation of spatial attention, thus amplifying the effects of attention and leading to increased alpha power lateralization. Indeed, such an anchoring effect may provide a productive perspective on prior demonstrations of object-based attention (Egly, Driver, & Rafal, 1994). While future work is needed to investigate the complex interrelationship between lateralized alpha power and online object representations, the present work clearly suggests that lateralized alpha power does not directly generate, and is dissociable from, the CDA.

### Conclusions

A growing body of evidence has shown that CDA and alpha power are tightly linked with the maintenance of information in the focus of attention. Here, we present new evidence that these two neural signals represent distinct facets of this online system. A topographic distribution of alpha power indexes the current locus of spatial attention, a process that is integral to both visual selection and the voluntary storage of items in WM. By contrast, CDA tracks the active maintenance of object files, the item-based representations that allow observers to integrate the ensemble of features and labels that are associated with visual objects. The dissociable activity of the CDA and alpha power suggests that the focus of attention is composed of at least two distinct but complementary neural processes, a conclusion with strong implications for both cognitive and neural models of this online storage system.

## Supplemental Materials

While the main manuscript focuses on the aggregate analysis of Experiments 1 and 2 to streamline the central message of the paper, here, we provide a more detailed analysis of the individual experiments.

### Rationale for Collapsing Experiment 1a and 1b

In a preliminary analysis, we examined whether the small variations in the task design between Experiments 1a and 1b had an effect on the observed results. For this purpose, we ran repeated measures ANOVAs for each analysis (e.g., Behavior, CDA, etc.) with the within-subjects factors Task (Working Memory [WM], Attention) and Set Size (2, 4 items) and the between-subjects factor Experiment (1a, 1b). With one exception (described next) there was no main effect of Experiment and no interaction of Experiment with any other factor, *p* ≥ .49. For the CDA analysis, there was a significant interaction of Laterality, Task, and Experiment, F(46) = 4.28, *p* = .04, η_p_^2^ = .09. To explore whether task lateralization varied across experiments, we ran a repeated measures ANOVA with the factors Laterality (Contralateral [“contra”], Ipsilateral [“ipsi”]) and Task (WM, Attention) for Experiment 1a and 1b separately. For both experiments, there was a significant interaction of Laterality and Task, *p* < .001. Specifically, there was a larger difference in laterality between the two tasks for Experiment 1a (M = −.50, SD = .33) than for Experiment 1b (M = −.31, SD = .31). The triple interaction between Laterality, Task and Experiment was not important for our central question which focused on the cognitive processes that yield lateralized activity. Thus, the data were collapsed across Experiments 1a and 1b.

## Experiment 1

### Behavior Results

#### Working memory

WM performance (Figure S1) was converted to a capacity score, K, calculated as K = N x (H-FA). N is the set-size; H is the hit rate; and FA is the false alarm rate (Cowan, 2001). We only analyzed “target absent” trials, as we excluded “target present” trials (10% of total trials) from the CDA and alpha analyses. For the WM trials, we compared performance between set size 2 and set size 4 using a two-tailed, paired-samples t-test. There was a significant difference in K score between set size 2 and 4, t(47) = −3.14, *p* = .003. Although the effect was smaller than usual, participants remembered significantly more items on set size 4 trials (M = 1.73, SD = .77) than set size 2 trials (M = 1.52, SD = .40).

#### Attention

Accuracy (Figure S1) was essentially at ceiling for detecting whether a line was present (Set Size 2: M = .99, SD = .01; Set Size 4: M = .97, SD = .02). On the rare trials in which lines were presented, participants correctly reported the orientation of the target line more frequently on set size 2 (M = .82, SD = .13) trials than on Set Size 4 (M = .78, SD = .15) trials, t(47) = 2.56, *p* = .01. Thus, monitoring four locations was more difficult than monitoring two locations.

**Figure S1.**
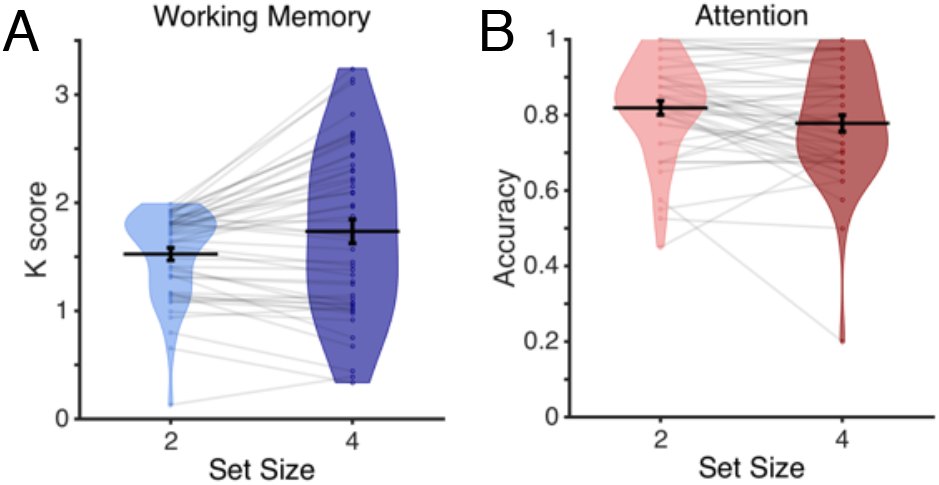
Behavioral performance for Experiment 1. (A) Average K score in the WM task for each experiment. The distribution of K scores for all participants is represented by the violin plot. Dots and light gray lines represent individual participants, performance. (B) Average accuracy in the attention task when participants had to discriminate the orientation of the target line.

### Contralateral Delay Activity Results

Recall that the WM condition required participants to direct attention to one side and store the colors of the objects in WM, while the attention task required only the deployment of spatial attention without storage of the objects in the sample array. As shown in Figure S2, a significant CDA was observed only in the WM condition.

To characterize the apparent differences in CDA amplitude across task, we ran a 2×2×2 repeated measures ANOVA with the factors Laterality (contra, ipsi), Task (WM, attention), and Set Size (2, 4 items) on data averaged from 400 to 1450 ms after stimulus onset. In this analysis, a significant effect of Laterality (i.e., greater negativity contralateral than ipsilateral to the sample array) provides evidence for a reliable CDA. This analysis revealed a significant main effect of Laterality, *F*(1,47) = 36.26, *p* < .001, η_p_^2^ = .44, and Set Size, *F*(1,47) = 12.85, *p* = .001, η_p_^2^ = .21. There was also a significant 2-way interaction of Laterality and Task, *p* < .001, and a significant three-way interaction of Laterality, Task, and Set Size, *F*(1,47) = 4.96, *p* = .03, η_p_^2^ = .10. No other effects were significant, *p* ≥ .60.

To characterize the significant interactions, we ran separate repeated measures ANOVAs for the WM and Attention tasks, with the within-subjects factors Laterality (contra, ipsi) and Set Size (2, 4 items). For the WM task, there was a significant main effect of Set Size, *F*(1,47)= 6.56, *p* = .01, η_p_^2^ = .12, and of Laterality, *F*(1,47) = 66.59, *p* < .001, η_p_^2^ = .59. The main effect of Set Size does not provide information about the CDA component, because the effect was found in data collapsed across Laterality. The main effect of Laterality, however, demonstrates that there was a reliable CDA in the WM condition. For the Attention task, the repeated measures ANOVA revealed a significant main effect of Set Size, *F*(1,47) = 10.07, *p* = .003, η_p_^2^ = .18. Again, this effect is not diagnostic regarding the CDA, because the data were collapsed across laterality. Critically, there was no main effect of Laterality in the Attention task, *F*(1,47) = .95, *p* = .33, η_p_^2^ = .02, suggesting that deploying covert attention to the locations of the sample items was not sufficient to drive the CDA.

Finally, the three-way interaction between Laterality, Set Size, and Task reflects the finding that set size effects in the attention task went modestly in the opposite direction of a typical CDA, with greater negativity in the set size 2 condition (M = −.07, SD = .29) than in the set size 4 condition (M = .001, SD = .29). This was not the case for the WM condition. One unusual finding was that we did not observe a significant interaction between Set Size and Laterality on the CDA amplitude in the WM task, despite many past demonstrations that CDA amplitude is sensitive to changes in mnemonic load (Luria, Balaban, Awh, & Vogel, 2016). CDA amplitude was numerically higher with the larger set size, but this was not statistically reliable. We think this may be due to unusually low WM performance for this group of subjects. While participants stored significantly more items in the set size 4 condition compared to the set size 2 condition, this difference was quite small (about 0.1 items); this modest difference in the number of items stored in the two conditions probably explains why a reliable change in CDA amplitude was not observed. Nevertheless, Experiment 2 will replicate the core empirical pattern while revealing a clear effect of WM load on CDA amplitude. Furthermore, aggregate analysis of CDA amplitude across Experiments 1 and 2 will also show a robust set size effect in line with past findings in the literature (see the main text).

To summarize the CDA analysis for Experiment 1, the pattern of the CDA was strikingly different for the WM and Attention tasks. When participants were instructed to encode and maintain object representations, we observed a robust CDA. However, when participants were instructed to deploy covert attention to the locations of the same squares to perform a demanding target discrimination task, we saw no evidence of the CDA.

**Figure S2.**
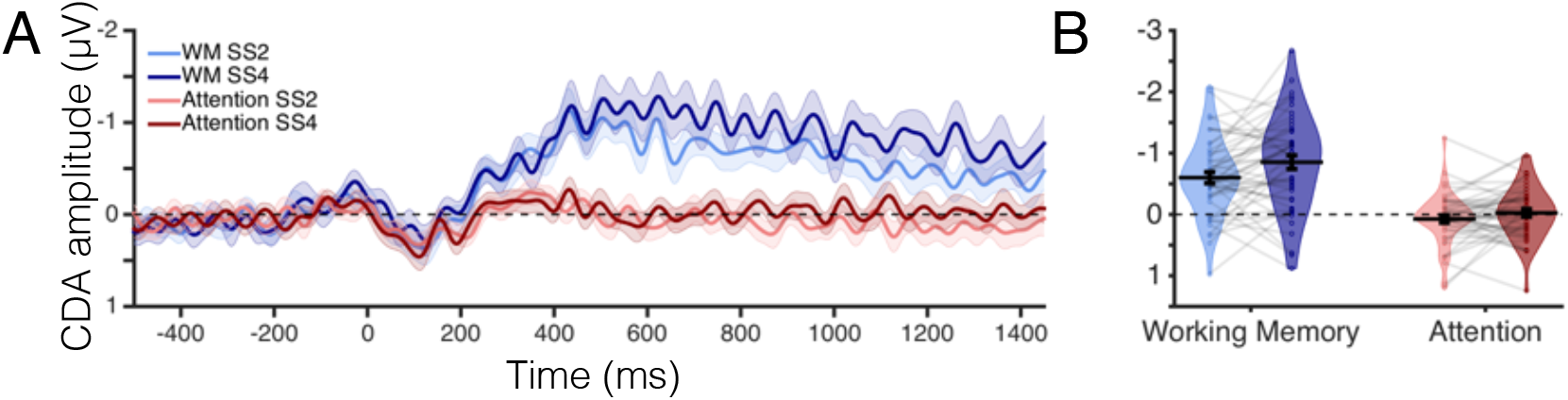
Contralateral delay activity for Experiment 1. (A) CDA amplitude (μV) over time. Timepoint zero marks the onset of the memory array, and timepoint 1450 marks the onset of the response array. (B) Average CDA amplitude (μV) during the time window of interest, 400 to 1450 ms. The distribution of CDA amplitudes for all participants is represented by the violin plot. Dots and light gray lines represent individual participants.

### Lateralized Alpha Power Results

As Figure S3 shows, we observed the typical suppression of alpha power contralateral to the relevant hemifield in both tasks, though it was larger in the WM task. We confirmed these impressions with a repeated measures ANOVA on the average alpha power in the same window as the CDA with the factors Laterality (contra, ipsi), Task (attention, WM) and Set Size (2, 4 items). The analysis revealed a main effect of Laterality, *F*(1,47) = 22.99, *p* < .001, η_p_^2^= .33, and a significant interaction of Laterality and Task, *F*(1,47) = 15.28, *p* < .001, η_p_^2^ = .25. All other effects were not significant, *p* ≥ .23. Mean alpha lateralization was stronger in the WM (M = −.46, SD = .45) than in the Attention (M = −.04, SD = .29) condition. Nevertheless, follow-up two-way paired-samples t-tests revealed that alpha power was significantly lateralized in all conditions, WM set size 2: *t*(47) = −5.25, *p* < .001; WM set size 4: *t*(47) = −4.91, *p* < .001; Attention set size 2: *t*(47) = −3.97, *p* < .001; Attention set size 4: *t*(47) = −3.29, *p* = .002. Thus, in both the WM and Attention conditions, we saw clear evidence of reduced alpha power in posterior contralateral electrodes, consistent with the hypothesis that both the WM and Attention tasks recruited covert spatial attention to the locations of the sample items. However, we also observed an effect of task, with more robust lateralization of alpha power in the WM task.

**Figure S3.**
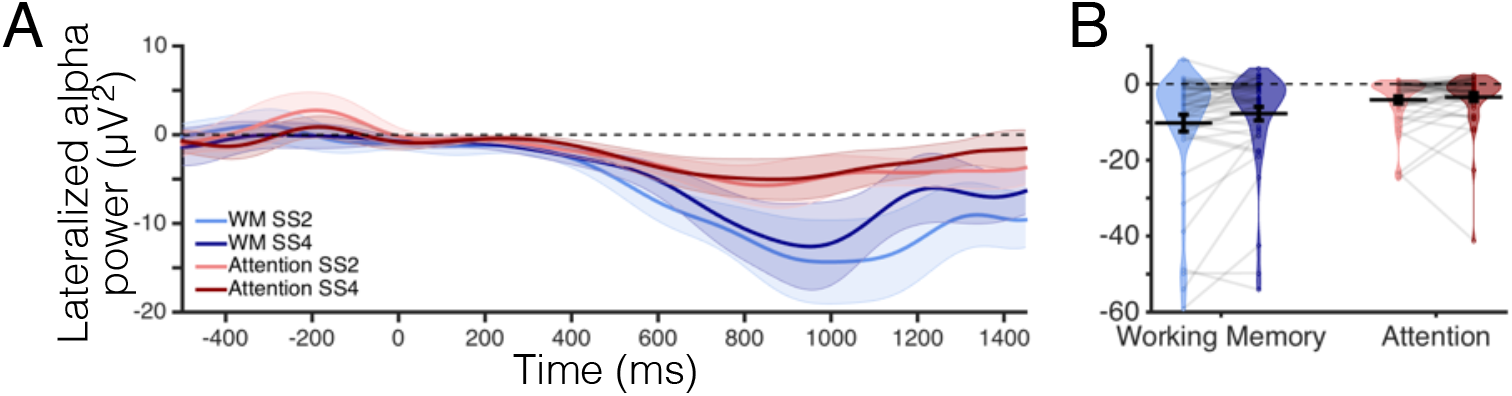
Lateralized alpha power for Experiment 1. (A) Lateralized alpha power (μV^2^) over time. Timepoint zero marks the onset of the memory array, and timepoint 1450 marks the onset of the response array. (B) Average lateralized alpha power (μV^2^) during the time window of interest, 400 to 1450 ms. The distribution of lateralized alpha power for all participants is represented by the violin plot. Dots and light gray lines represent one participant’s alpha power.

### Pupil Dilation Results

To compare pupil dilation (Figure S4) between the WM and Attention tasks collapsed across set size, we ran a two-way paired-samples t-test on the averaged pupil dilation from the same time window as the CDA. We did this separately for each task. There was no significant difference in average pupil dilation between the WM (M = .65, SD = 2.00) and Attention (M = .86, SD = 1.78) tasks, *t*(43)= −.83, *p* = .41. Thus, pupil dilation provided no indication that stronger lateralized activity in the WM task was due to increased effort. Indeed, an aggregate analysis below will provide evidence that this correlate of cognitive effort was actually higher in the Attention task.

**Figure S4.**
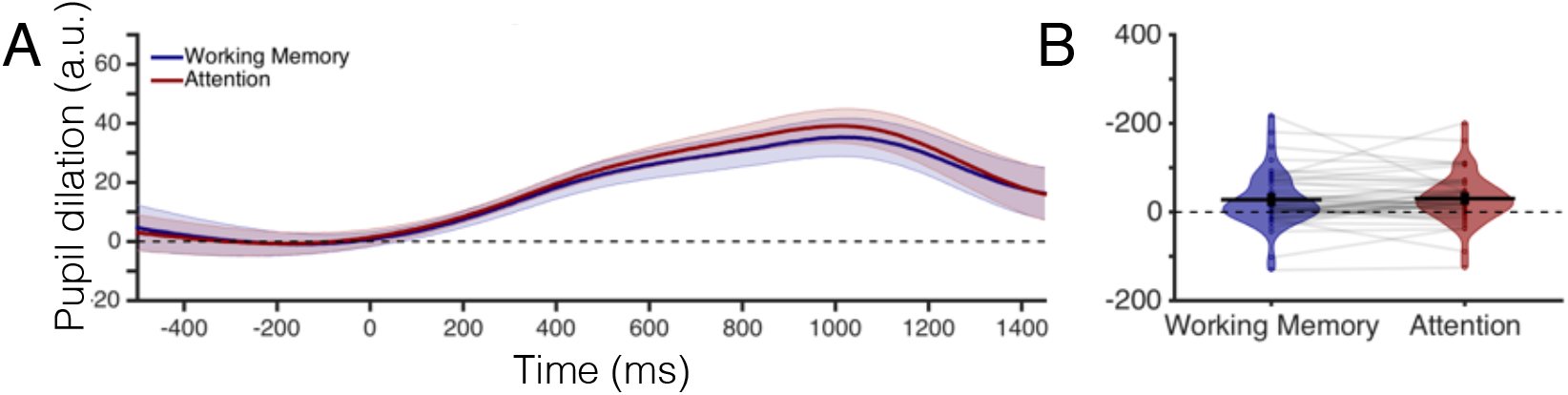
Task-evoked pupil response for Experiment 1. (A) Pupil dilation (arbitrary units) over time. Timepoint zero marks the onset of the memory array, and timepoint 1450 marks the onset of the response array. (B) Average pupil dilation (arbitrary units) during the time window of interest, 400 to 1450 ms. The distribution of pupil dilation for all participants is represented by the violin plot. Dots and light gray lines represent one participant’s pupil dilation.

### Horizontal Gaze Position Results

To compare horizontal gaze position between the tasks (during the same time window used to measure the CDA) we ran a 2×2 repeated measures ANOVA on horizontal eye position with the factors Task (WM, Attention) and Set Size (2, 4 items). This analysis revealed that there was no main effect of Task or Set Size, and no significant interaction between these two factors, *p* ≥ .12 for all effects. Thus, participants moved their eyes the same amount in all conditions. In all conditions, participants moved their eyes less than 0.017 degrees of visual angle, which is smaller than the size of the fixation dot.

## Experiment 2

### Rationale for Collapsing Experiment 2a and 2b

In a preliminary analysis, we examined whether the small variations in task design between Experiments 2a and 2b had an effect on the observed results. For this purpose, we ran repeated measures ANOVAs for each analysis (e.g., Behavior, CDA, etc.) with the within-subjects factors Task (WM, Attention) and Set Size (2, 4 items) and the between-subjects factor Experiment (2a, 2b). There was no main effect of Experiment for any of the analyses, *p*>=.10, and no interaction of Experiment with any other factor (*p* > .10). Therefore, subsequent analyses collapsed the data across 2a and 2b.

### Behavior Results

We separately analyzed performance for the WM and attention tasks (Figure S5). For the WM task, there was a significant difference in K score between Set Size 2 and 4, *t*(48)= −4.1, *p* < .001. Participants remembered significantly more items on set size 4 (M = 1.43, SD = .69) than set size 2 trials (M = 1.23, SD = .49).

In the Attention task, participants had a high rate of detecting whether a line was present (Set Size 2: M = .97, SD = .02; Set Size 4: M = .96, SD = .02). To compare performance between set size 2 and 4 when participants had to discriminate the orientation of the target line, we used a two-tailed, paired-samples t-test. Participants correctly reported the orientation of the target line more frequently on Set Size 2 (M = .75, SD = .10) than on Set Size 4 (M = .69, SD = .11) trials, *t*(48) = 4.00, *p* < .001. Thus, monitoring four locations was more difficult than monitoring two locations.

**Figure S5.**
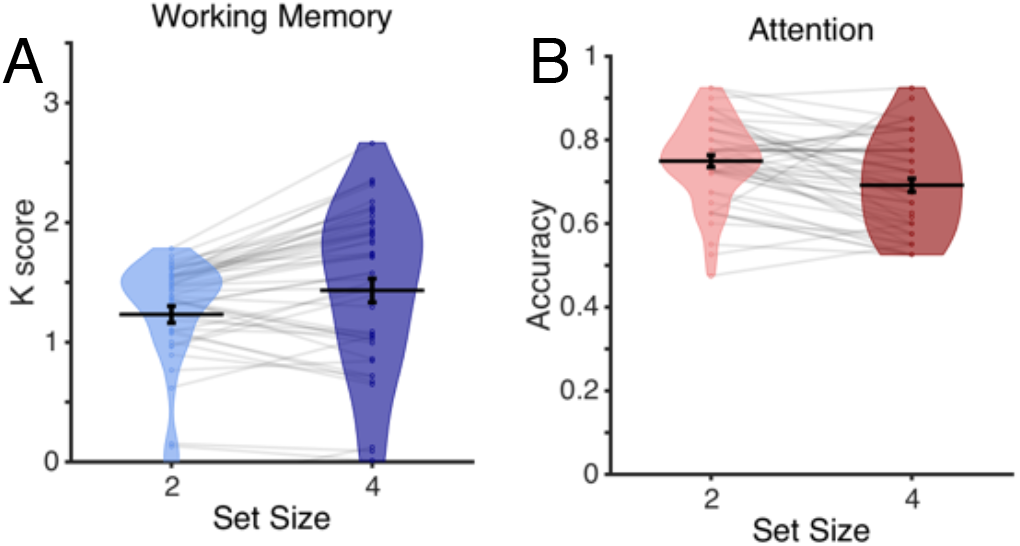
Behavioral performance for Experiment 2. (A) Average K score in the WM task for each experiment. The distribution of K scores for all participants is represented by the violin plot. Dots and light gray lines represent one participant’s performance. (B) Average accuracy in the attention task when participants had to discriminate the orientation of the target line.

### Contralateral Delay Activity Results

As shown in Figure S6, CDA amplitude was different across the WM and Attention tasks, with more robust lateralized activity and effects of set size in the WM task. This impression was confirmed with a repeated measures ANOVA with the factors Task (WM, Attention), Laterality (contra, ipsi) and Set Size (2, 4 items). The repeated measures ANOVA revealed significant main effects of Laterality, *F*(1,48) = 39.38, *p* < .001, η_p_^2^ =.45, and Set Size, *F*(1,48) = 21.03, *p* < .001, η_p_^2^ = .31. Critically, there was a reliable interaction between Laterality and Task, *F*(1,48) = 23.68, *p* < .001, η_p_^2^ = .33, showing that CDA amplitude differed between the WM and Attention tasks. There were also significant interactions between Task and Set Size, *F*(1,48) = 5.72, *p* = .02, η_p_^2^ = .11, and between Laterality and Set Size *F*(1,48) = 16.88, *p* < .001, η_p_^2^ = .26; because these interactions refer to data collapsed across Laterality or Task, they are not informative regarding the central question of which task requirements generate the CDA. Finally, given our hypothesis that the CDA may be specifically tied to the formation of object representations, we also predicted a triple interaction between Laterality, Task and Set Size, because load-dependent CDA effects should be more pronounced in the WM task. This interaction was trending but did not reach conventional thresholds for significance, *F*(1,48) = 3.72, *p* = .06, η_p_^2^ = .07. We re-visit this question below with a more sensitive aggregate analysis of Experiments 1 and 2 together.

To further delineate the key interaction between Laterality and Task, we ran a follow-up paired-samples t-tests, collapsed across set size. Examining the difference between contralateral and ipsilateral activity in each task, we observed reliable effects of Laterality for both the WM and Attention tasks, *p* ≤ .03 for all conditions. Critically, however, the effect of Laterality was substantially larger in the WM task, (M = −.49, SD = .50) than in the Attention task (M = −.17, SD = .35), t(48) = −4.88, *p* < .001. Thus, Experiment 2 replicated the broad empirical pattern in Experiment 1; the CDA was far stronger when participants were instructed to store object representations in WM than when they were instructed to attend those locations in anticipation of upcoming targets.

The marked difference in CDA amplitude between the WM and Attention tasks is particularly striking in light of their strong similarity. Indeed, while the labels for the WM and Attention tasks highlight their respective storage and selection requirements, both tasks actually require the sustained maintenance of spatial information. In the WM task, participants held the positions of the sample items in mind to facilitate change detection, while in the Attention task, participants held a sustained focus of spatial attention at specific positions. Nevertheless, CDA amplitude was more than twice as high in the WM task, motivating the conclusion that the CDA is tied to object representations held in WM *per se*, and not simply the deployment of covert attention – even when an attention task requires the maintenance of spatial information. If this is correct, however, it raises the question of why we saw reliable a CDA in the Attention task. Our working hypothesis is that this modest effect of Laterality could reflect the occasional storage of object representations during the Attention task, but the current study does not provide a clear way to test this possibility. Nevertheless, while this question can’t be fully answered, Experiment 2 did replicate the striking divergence in CDA amplitude between the WM and Attention tasks, in line with the hypothesis that this neural signal may be specifically tied to item storage in visual WM.

**Figure S6.**
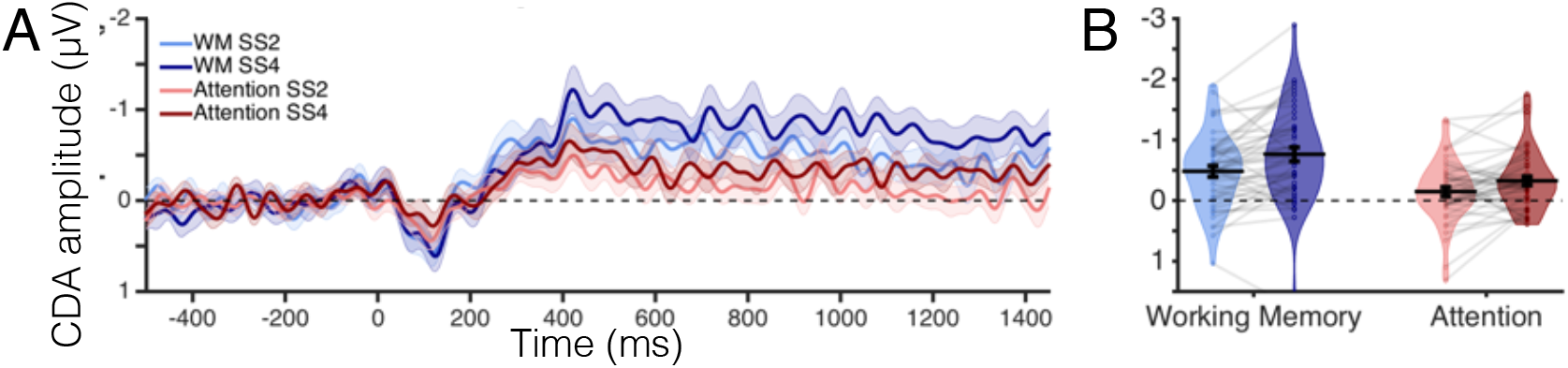
Contralateral delay activity for Experiment 2. (A) CDA amplitude (μV) over time. Timepoint zero marks the onset of the memory array, and timepoint 1450 marks the onset of the response array. (B) Average CDA amplitude (μV) during the time window of interest, 400 to 1450 ms. The distribution of CDA amplitudes for all participants is represented by the violin plot. Dots and light gray lines represent one participant’s CDA amplitude.

### Lateralized Alpha Power Results

We observed the typical suppression of alpha power contralateral to the relevant hemifield in both tasks (Figure S7). Once again, this effect was larger in the WM task. We confirmed these impressions with a repeated measures ANOVA on average alpha power with the factors Laterality (contra, ipsi), Task (attention, WM) and Set Size (2, 4 items). The analysis revealed a significant main effect of Laterality, *F*(1,48) = 24.17, *p* < .001, η_p_^2^ = .34, and a significant interaction between Laterality and Task, *F*(1,48) = 13.38, *p* < .001, η_p_^2^ = .22. Follow-up two-way paired samples t-tests on the difference between contralateral and ipsilateral activity for each condition revealed significant alpha power suppression in all conditions, *p* ≤ .001 for both set sizes in both the WM and Attention tasks. No other effects were significant, *p* ≥ .10.

To characterize the interaction of Laterality and Task, we collapsed Laterality into a difference wave (contra minus ipsi). We then compared this difference wave for the Attention and WM conditions, averaged across Set Size, with a two-way paired samples t-test. This analysis revealed a significant difference in alpha power lateralization between the WM and Attention tasks, *t*(48) = −3.66, *p* = .001. Alpha power was more suppressed in contralateral relative to ipsilateral electrodes during the WM (M = −9.16, SD = 13.38) than during the Attention (M = −4.01, SD = 6.73) task.

To summarize, in both tasks, we found clear evidence of sustained covert orienting towards the relevant hemifield. This makes the important point that the lack of a robust CDA in the Attention task is not due to a failure to maintain attention towards the relevant hemifield. In addition, replicating the finding from Experiment 1, we observed reliably stronger lateralization of alpha power when participants were instructed to store the sample items in WM. Thus, both neural signals suggest a distinction between the maintenance of items in working memory, and the maintenance of spatial attention at the position of those items.

**Figure S7.**
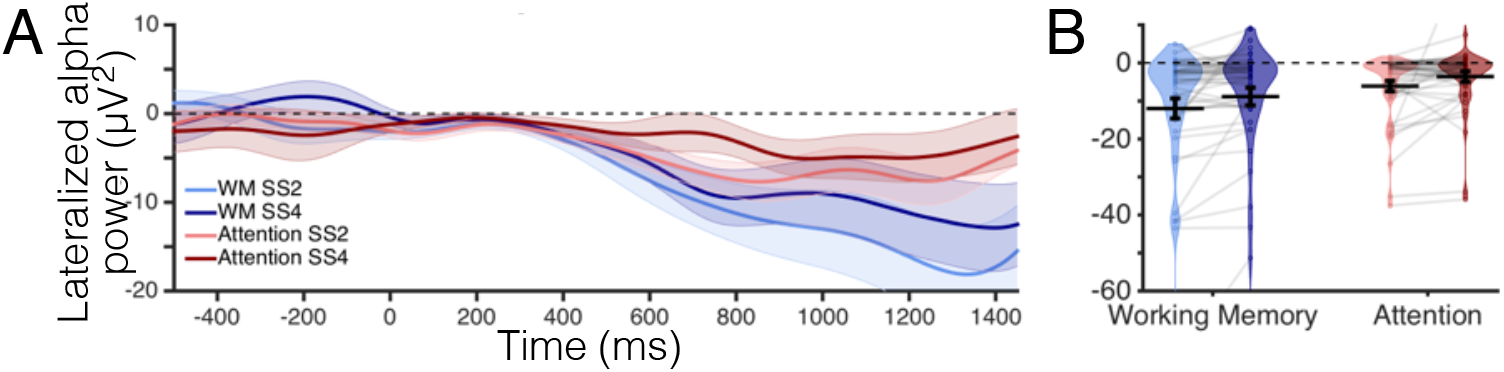
Lateralized alpha power for Experiment 2. (A) Lateralized alpha power (μV^2^) over time. Timepoint zero marks the onset of the memory array, and timepoint 1450 marks the onset of the response array. (B) Average lateralized alpha power (μV^2^) during the time window of interest, 400 to 1450 ms. The distribution of lateralized alpha power for all participants is represented by the violin plot. Dots and light gray lines represent one participant’s alpha power.

### Pupil Dilation Results

To compare the task-evoked pupil response (Figure S8) between the WM and Attention tasks collapsed across set size, we ran a two-way paired-samples t-test. This analysis revealed greater task-evoked pupil dilation in the Attention task (M = .60, SD = .29) than in the WM task (M = −.07, SD = 2.36), *t*(43) = −3.65, *p* = .001, suggesting that the Attention task elicited greater cognitive effort. Therefore, the robust CDA in the WM condition is unlikely to reflect greater cognitive effort in the WM than the Attention task.

**Figure S8.**
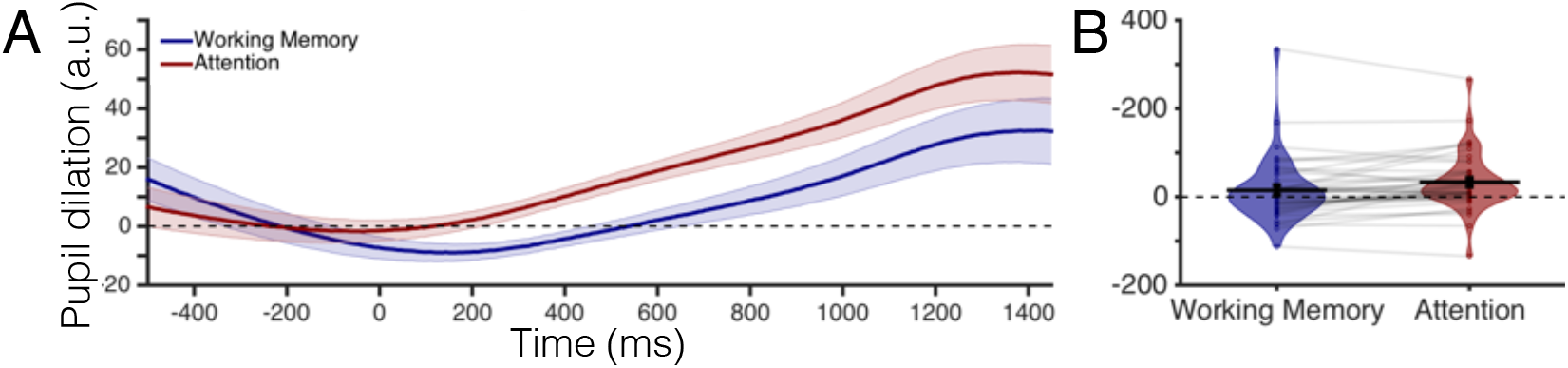
Task-evoked pupil response for Experiment 2. (A) Pupil dilation (arbitrary units) over time. Timepoint zero marks the onset of the memory array, and timepoint 1450 marks the onset of the response array. (B) Average pupil dilation (arbitrary units) during the time window of interest, 400 to 1450 ms. The distribution of pupil dilation for all participants is represented by the violin plot. Dots and light gray lines represent one participant’s pupil dilation.

### Horizontal Gaze Position Results

To determine whether horizontal gaze position varied across tasks, we ran a 2×2 repeated measures ANOVA with the factors Task (WM, Attention) and Set Size (2, 4 items). This analysis revealed a significant main effect of Task, *F*(1,42) = 4.78, *p* = .03, η_p_^2^= .10. Participants moved their eyes more during the WM (M = .03, SD = .03) than the Attention (M = .02, SD = .02) task. We would like to point out that the difference in eye movements between these two tasks is only about .01 degrees, which is smaller than the diameter of the fixation point. Nevertheless, to determine whether these differences in horizontal gaze position drive the differences in the CDA that we observe, we compared horizontal gaze position during the time window when the CDA initially emerges, 400 to 925 ms after stimulus onset. The 2×2 repeated measures ANOVA with factors Task (WM, Attention) and Set Size (2, 4 items) for the average horizontal gaze position during this time window revealed that there was no significant main effect of Task or Set Size and no interaction of these two factors, *p* ≥ .06 for all effects. This indicates that during the time window when the CDA is ramping up, participants moved their eyes the same amount in all conditions. Therefore, any differences in the CDA that we observe are not driven by differences in horizontal eye movements. As a follow-up analysis we also analyzed horizontal eye movement averaged over the end of the trial, 926 to 1450 ms, with another 2×2 ANOVA with factors Task (WM, Attention) and Set Size (2, 4 items). This analysis revealed that differences in horizontal eye movements between the WM and Attention tasks emerged toward the end of the trial, significant main effect of Task, *F*(1,43) = 4.40, *p* = .04, η_p_^2^ = .09. All other effects and interactions were not significant, *p* ≧ .06.

**Contributions:** All authors conceived and designed the experiments. N.H. and K.A. collected the data, performed analyses, and drafted the manuscript; N.H., K.A., E.A., and E.V. revised the manuscript.

